# Coexistence of bacteria with a competition-colonization tradeoff on a dynamic coral host

**DOI:** 10.1101/2024.09.15.612558

**Authors:** Theo L. Gibbs, Kyle J.-M. Dahlin, Joe Brennan, Cynthia B. Silveira, Lisa C. McManus

**Affiliations:** Lewis-Sigler Institute for Integrative Genomics, Princeton University, Princeton, New Jersey 08544 USA; Department of Mathematics and the Center for the Mathematics of Biosystems, Virginia Tech, Blacksburg, Virginia 24061 USA; Center for Population Biology, University of California, Davis, California 95616 USA; Department of Biology, University of Miami, Coral Gables, Florida 33146 USA; Hawai’i Institute of Marine Biology, University of Hawai’i at Mānoa, Kāne’ohe, Hawaii 96744 USA

**Author notes:** Equal contribution (Cynthia B. Silveira, Lisa C. McManus).

**Keywords:** Holobiont, competition-colonization, corals, microbial dynamics

## Abstract

Many macroscopic organisms enter into tightly linked symbiosis with microbial communities. Although experimental work has demonstrated the importance of these symbioses, a theoretical understanding of stable, multi-scale coexistence remains underdeveloped. Here, we explored how the competition-colonization tradeoff, a classic coexistence mechanism, operates when bacterial species compete for a dynamic biological host. Specifically, we introduce a model where corals are colonized by fast-growing mutualists and slow-growing pathogens. We found that the vital rates of the host coral influenced coexistence outcomes between bacterial types. Notably, pathogen-induced host death expanded the region of parameter space where coexistence was stable for all three species and mutualistic bacteria enabled coexistence in systems that would have otherwise collapsed. These findings provide new insights into the interplay between microbial interactions and macroscopic processes. Our work illustrates how host-microbe interactions can shape ecosystem stability, providing a theoretical framework applicable to a wide range of symbiotic systems.

## Introduction

Across the tree of life, microbial communities and macroscopic host organisms enter into closely intertwined symbiotic relationships. The majority of plant species rely on mycorrhizal fungi to acquire mineral nutrients, and in turn, these plants provide organic nutrients to the fungi (Jansa et al., 2008, 2011; Smith and Read, 2010). The gut microbiome affects many aspects of human health, and reciprocally, these microbial populations require the nutrients and habitat provided by their hosts (Eloe-Fadrosh and Rasko, 2013). Corals provide habitat for photosynthetic zooxanthellae and receive oxygen, organic carbon and other compounds in return (Roth, 2014). These examples demonstrate that microscopic and macroscopic organisms often form integrated mutualisms critical to the functioning of both parties, leading to the development of the ‘holobiont’ concept to describe such closely intertwined symbiotic systems (Margulis, 1992; Rohwer et al., 2002).

This integrative approach is critical to understanding the ecology and evolution of macroscopic hosts. Microbiota have been shown to modify the behavior and social interactions among hosts, and can even produce reproductive isolation, potentially inducing speciation (Bell and Bordenstein, 2022). Despite these insights, there has been comparatively little progress on the population theory of holobionts, particularly in systems with nested hierarchies of organisms. We have a quantitative understanding of the conditions under which microbial species coexist (Gause and Witt, 1935; MacArthur, 1970; Monod, 1949; Chase and Leibold, 2003; Huisman and Weissing, 1999; Posfai et al., 2017; Marsland III et al., 2019; Butler and O’Dwyer, 2018), theory for how mutualistic interactions between organisms shape community composition (Bronstein, 2015, 2001; Valdovinos et al., 2013; Valdovinos, 2019; Martignoni et al., 2020), and mathematical models describing the population biology of host species (Hastings, 1996), but these bodies of work have not been adequately merged (see Roughgarden 2023 for notable recent progress). Developing a comprehensive population theory for holobionts is crucial for understanding the dynamics of these complex symbiotic systems.

In our exploration of holobiont dynamics, we leverage the classic competition-colonization trade-off framework, a fundamental concept in theoretical ecology (Levins and Culver, 1971; Levin and Paine, 1974; Horn and MacArthur, 1972; Hastings, 1980; Tilman, 1994). This model posits that species cannot excel at both competing for resources and colonizing new habitats simultaneously. Superior competitors can dominate established habitats but may struggle to disperse effectively, while strong colonizers can quickly occupy new or disturbed areas but are out-competed in stable environments (Levins and Culver, 1971; Yu and Wilson, 2001). This trade-off has been widely observed across various ecosystems, from insects to microbial populations, and helps explain how diverse species can coexist in spatially structured environments (Cadotte et al., 2006; Cadotte, 2007; Livingston et al., 2012; Stanton et al., 2002; Wetherington et al., 2022). Recent research has shown that bacteria within the coral microbiome engage in a similar competition-colonization dynamic (Wang et al., 2022), providing a compelling system for exploring these ecological principles in the context of host-microbe interactions.

Corals harbor a diverse community of bacteria, microeukaryotes, archaea, and viruses in their tissues and mucus (Knowlton and Rohwer, 2003). The microbiome is an integral component of coral health and, therefore, reef ecosystem functioning and response to environmental change (Voolstra et al., 2024). Putative mutualistic bacteria can help corals avoid pathogens and survive heat stress (Krediet et al., 2013; Santoro et al., 2021). On the other hand, temperature changes modulate microbial species interactions that can lead to coral death (Rubio-Portillo et al., 2020). To investigate the effects of inter-species interactions in the coral holobiont, we model pathogenic and mutualistic microbial species colonizing a coral colony following the classic competition-colonization trade-off but where the patch landscape is itself a host species undergoing its own dispersal dynamics. Although our analysis is general, we are particularly motivated by the findings from a set of recent experiments with the coral *Galaxea fascicularis* and two bacterial species isolated from its gastrointestinal cavity: *Vibrio coralliilyticus*, an opportunistic pathogen, and a commensal *V. alginolyticus* (Wang et al., 2022). Briefly, the coral hosts’ fast-growing mutualistic bacteria are killed by the competing pathogenic *V. coralliilyticus* via prophage induction, mirroring the classic competition-colonization tradeoff (Reshef et al., 2006; Paul, 2008; McIlroy et al., 2019).

In this holobiotic context, we address the following three specific questions: 1) Are the rules for coexistence different when the “number of sites” – namely, the amount of live coral on a reef – is a dynamic biological entity?; 2) How does host death induced by the pathogen modify coexistence outcomes?; and 3) Can increased host growth due to colonization by a mutualist bacteria rescue host populations that would otherwise collapse? By addressing these questions, we aim to advance our understanding of the complex interactions within holobionts and their implications for ecosystem dynamics and species coexistence in host-microbe systems.

## Methods

### Model formulation

We consider a finite landscape of patches or reefs which can be in four different states – empty, occupied by a coral host, occupied by a coral host with bacterial pathogens, or occupied by a coral host with bacterial mutualists. The coral host colonizes empty patches and dies at a constant rate. The pathogenic bacteria can induce coral death, while the mutualistic bacteria can increase the patch colonization rate of the coral. Both types of bacteria can colonize patches occupied by the coral host, but the pathogenic bacteria is the dominant competitor compared to the mutualist because it can colonize patches occupied by the mutualist, killing it in the process. A simple system of ordinary differential equations that captures these biological processes is given by

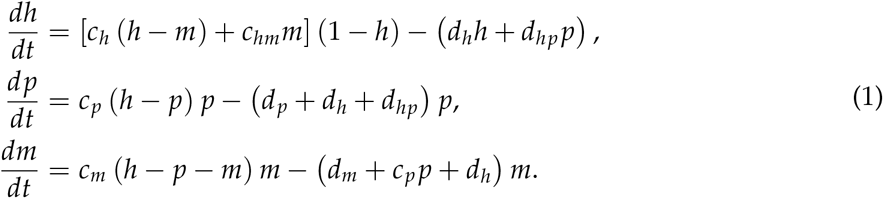

Here, *h* is the fraction of the landscape occupied by the coral host, *p* is the fraction of the landscape occupied by the pathogen and *m* is the fraction occupied by the mutualist. Both pathogens and mutualists must occupy a host individual to survive, so *p* + *m* ≤ *h*. Hosts that are not occupied by a mutualist colonize empty patches at rate *c*_*h*_, while hosts with a mutualist colonize empty patches at rate *c*_*hm*_. Under the assumption that patches are well-mixed across space, the 1 − *h* term represents the probability that a host disperses to an empty patch. Hosts die at an intrinsic rate *d*_*h*_ and an elevated rate *d*_*hp*_ when they are occupied by a pathogen. When individual hosts die, their associated bacteria do as well, so the *d*_*h*_ term enters into all of the equations, scaled by the proportion of coral occupied by the bacteria (*p* or *m*). Pathogens colonize host or mutualist patches at rate *c*_*p*_. Pathogens die at an intrinsic rate *d*_*p*_ and at a potentially elevated rate *d*_*hp*_ when they kill the host they are inhabiting. Mutualists colonize host patches at the rate *c*_*m*_. Mutualists experience natural mortality and additional mortality due to the competitive dominance of the pathogenic bacteria at rates *d*_*m*_ and *c*_*p*_ *p*, respectively. We present all the parameters of our model in Table A1. Because the life history traits corresponding to our model parameters are not commonly measured for coral-bacteria systems, we use a range of parameter values to explore the behavior of Equation (1).

### Numerical and theoretical analysis

To numerically integrate Equation (1), we used the Livermore Solver for Ordinary Differential Equations (LSODA) from the deSolve v1.25 package (Soetaert et al., 2010) in R version 3.6.1 (R Core Team, 2023). In addition to simulations of the dynamics, we also computed equilibria of Equation (1) algebraically and determined their stability analytically when it was tractable (see Appendix A). Following previous studies (Saavedra et al., 2017; Song et al., 2018), we call an equilibrium feasible when all species have positive abundances. Similarly, an equilibrium is stable when all the eigenvalues of the Jacobian associated with Equation (1) evaluated at the equilibrium have negative real parts. If an equilibrium is stable, then small perturbations away from the equilibrium densities asymptotically return to equilibrium. In the main text, we solve for feasible equilibria and determine their stability numerically. The code used to run simulations is available at https://github.com/theogibbs/HostCompCol.

## Results

We first delineate coexistence criteria when host traits impact the bacteria but neither the pathogen nor the mutualist impacts the host: the pathogen does not induce host death (*d*_*hp*_ = 0) and the mutualist does not benefit host colonization (*c*_*hm*_ = *c*_*h*_). Starting with parameters where the pathogen excludes the mutualist, increasing the host death rate first produces coexistence among the host, pathogen, and mutualist but eventually leads to pathogen exclusion (Fig. 1A). When the host does not die (*d*_*h*_ = 0), the host occupies all empty patches and we exactly recover the classic colonization tradeoff (leftmost panel of 1B and Levins and Culver (1971); Hastings (1980)). In this case, the mutualist must be a much faster colonizer than the pathogen to survive (Fig. 1B). As the host death rate increases, however, the coexistence region broadens because the mutualist is less easily excluded by the pathogen (Fig. 1B). While higher host death rates increase fraction of mutualists and decrease fraction of pathogens, higher host colonization rates have the opposite effect (Fig. 1C). In Appendix A, we derive a formula that explicitly show how the usual competition-colonization coexistence criteria are modified by host traits. **Our first key finding reveals that higher host densities favor the pathogen, while higher host death rates favor the mutualist when bacteria do not directly affect host mortality or growth**. That the bacterial species now compete for and colonize a dynamical biological entity rather than a static patch landscape leads to a departure from the standard coexistence results (e.g., the differences between the panels of Fig. 1B).

**Figure 1.**
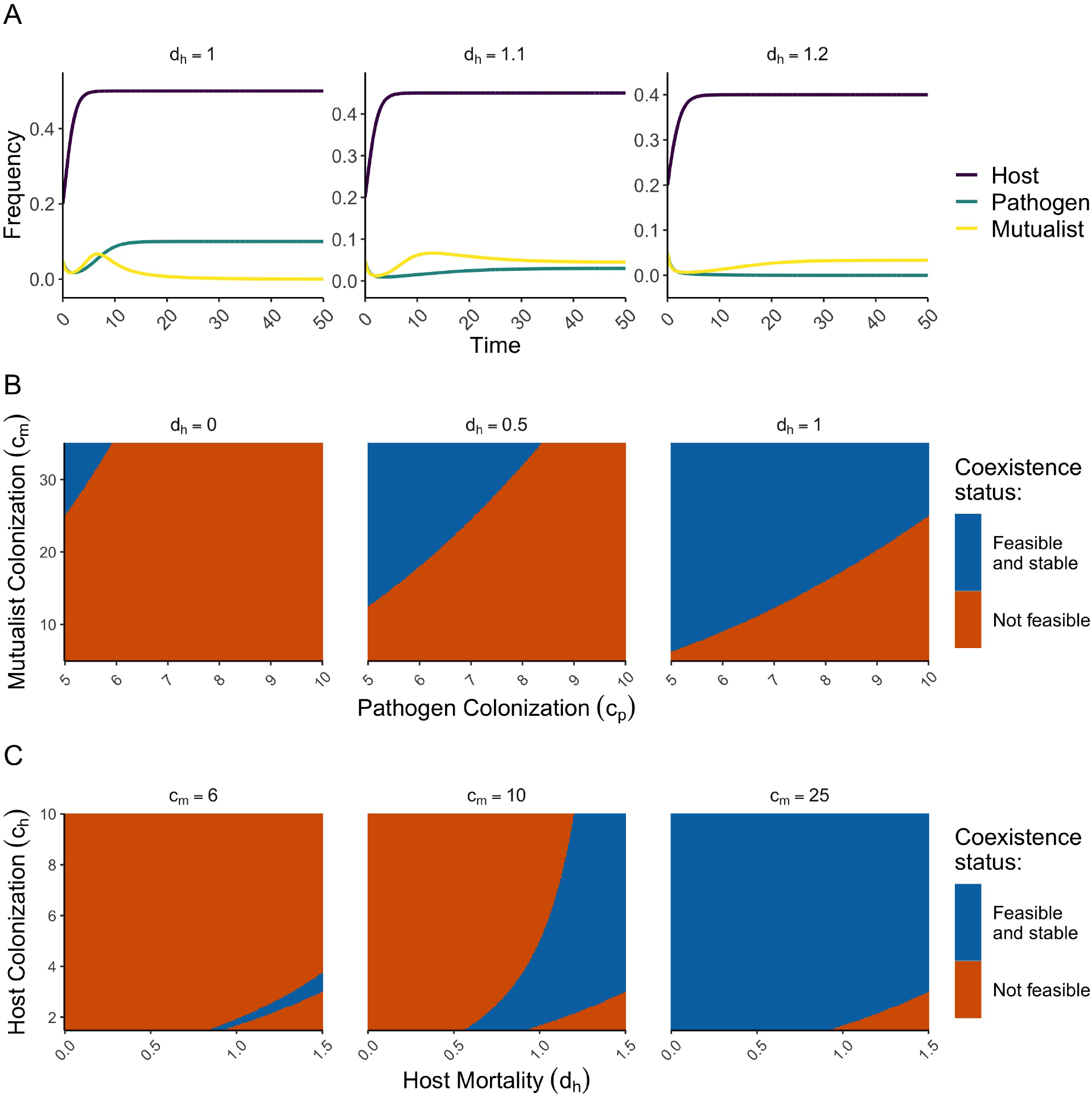
(A) Dynamics of the proportions of host, pathogen and mutualist over time for three choices of the intrinsic host death rate (panels). Here we assume that *d*_*hp*_ = 0 and *c*_*hm*_ = *c*_*h*_. (B) Heatmaps of when the equilibria identified in the main text are feasible and stable across a range of pathogen and mutualist colonization rates. Colors indicate whether or not the three species are predicted to stably coexist. Panels are different host death rates and the leftmost one in which *d*_*h*_ = 0 is the standard competition-colonization tradeoff. (C) Analogous to panel (B) except across a range of host mortality and colonization rates. Panels are different mutualist colonization rates. The remaining parameter values for panels B-C are given in Table A1.

Exploring pathogen-induced host death, we find that increasing *d*_*hp*_ allows for stable coexistence at lower mutualist colonization rates (*c*_*m*_) up to a point where the pathogen ceases to grow (Fig. 2B). Increasing *d*_*hp*_ also eliminates pathogen-occupied coral patches more quickly, eventually leading to higher densities of the mutualist and exclusion of the pathogen at equlibrium. Additionally, there is a resultant increase in the overall host death rate which, according to our first finding, favors the mutualist. Both of these mechanisms contribute to the stable coexistence of all three types. **Our second key finding is that pathogen-induced host death tends to favor coexistence, provided it is not so high as to preclude persistence of the pathogen.**

**Figure 2.**
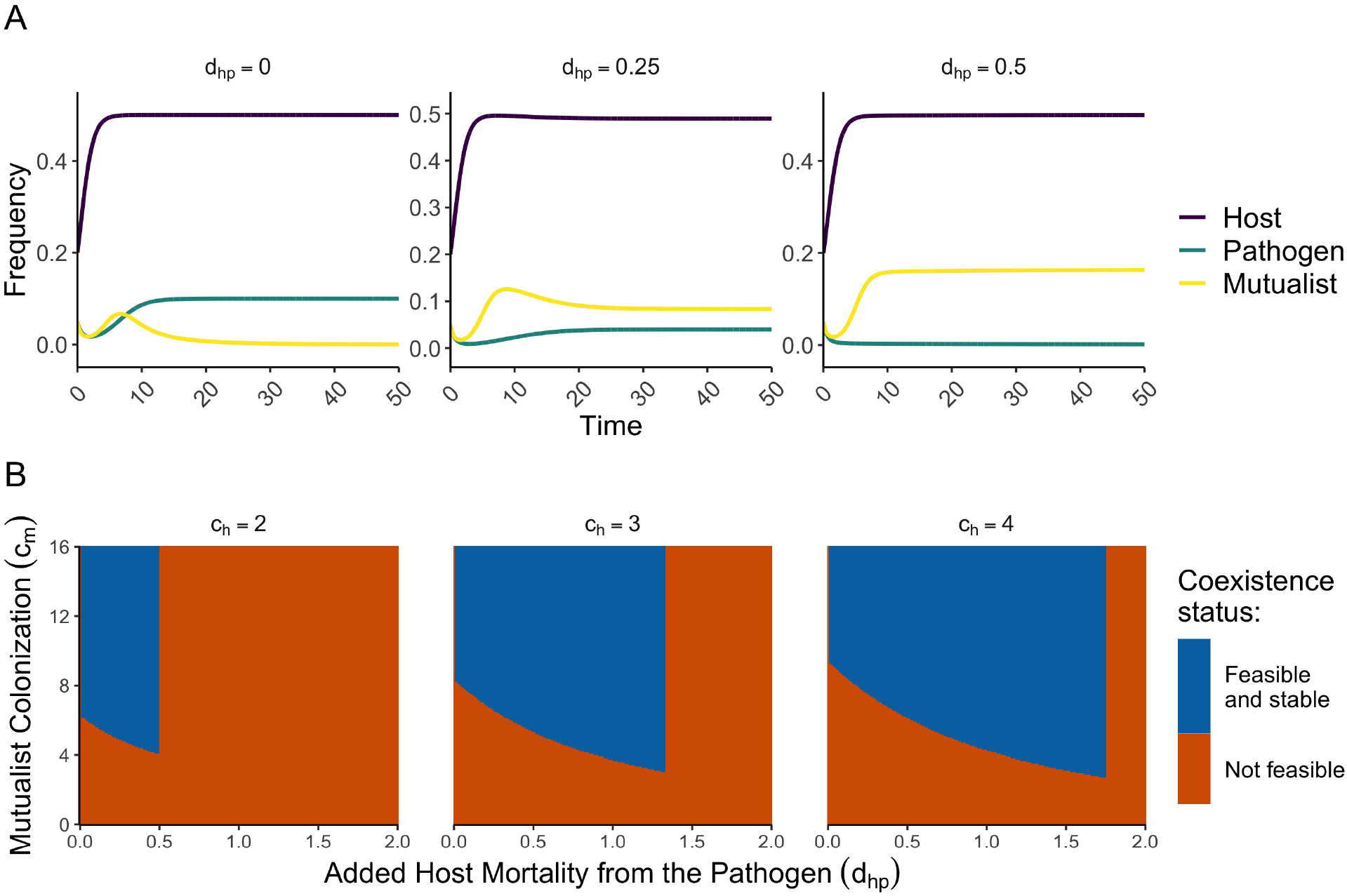
(A) Dynamics of the fractions of host, pathogen and mutualist over time for three choices of host mortality induced by the pathogen (panels). When the pathogen does not cause a larger death rate in the hosts, the mutualist is excluded, but at intermediate levels of additional mortality, the mutualist and pathogen coexist. At high levels of additional mortality, the pathogen is excluded. (B) Heatmaps of when the equilibria identified in the main text are feasible and stable across a range of pathogen-induced mortality and mutualist colonization rates. Colors indicate whether or not the three species are predicted to stably coexist. Here we assume that *c*_*hm*_ = *c*_*h*_. The remaining parameter values are given in Table A1.

To investigate mutualist-enhanced host growth effects, we modeled increased host colonization when occupied by the mutualist (*c*_*hm*_ > *c*_*h*_) without pathogen-induced host death (*d*_*hp*_ = 0). Even when the host cannot grow without the mutualist (*c*_*h*_ < *d*_*h*_), sufficiently high host colonization induced by the mutualist can rescue a system that would otherwise collapse, facilitating coexistence (Fig. 3A). For high intrinsic mutualist colonization rates (*c*_*m*_), coexistence occurs at a stable fixed point as in our previous results (top row of Fig. 3A). For low *c*_*m*_, however, the equilibrium where all species are present becomes unstable, and coexistence may be maintained through complicated dynamical behavior (Fig. 3A). Increasing the benefit of the mutualist to the host (*c*_*hm*_) shifts the system from collapse to dynamical coexistence to stable equilibrium co-existence (Fig. 3A-B). For hosts with higher intrinsic colonization rates that do not rely on the mutualists, the qualitative coexistence results remain the same (Fig. 3B). Surprisingly, as the added benefit of the mutualist to the host increases, the the fraction of hosts occupied by the mutualist decreases while the fraction of pathogen increases. However, this result aligns with our previous finding wherein the pathogen benefits more than the mutualist from any increase in the effective growth rate of the host.

**Figure 3.**
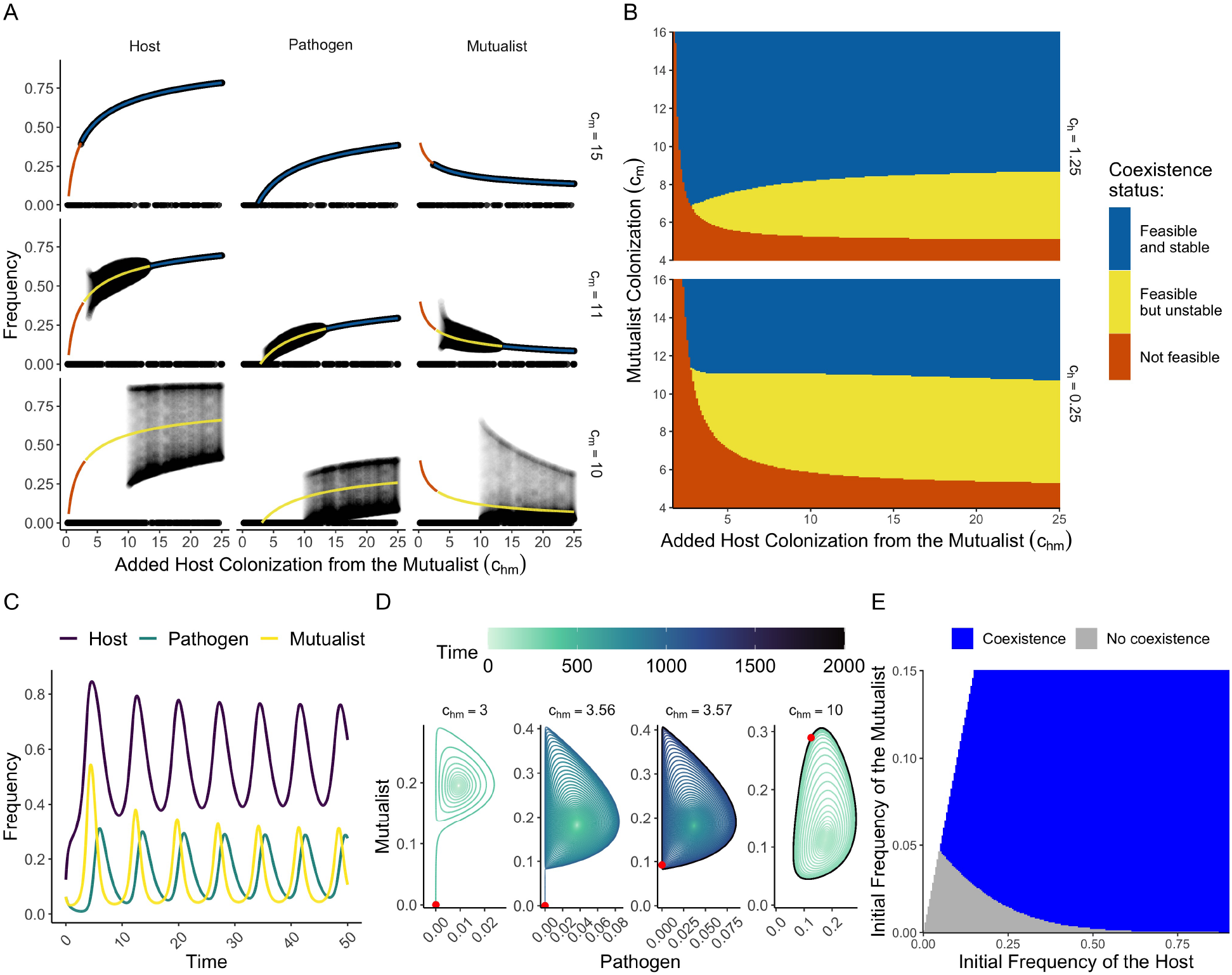
(A) Points are the fractions of the host, pathogen and mutualist (columns) in the last 250 timesteps of simulations starting from two different initial conditions across a range of additional host colonization rates caused by the mutualist. The rows show results for different rates of mutualist colonization. Lines show predicted equilibrium fractions and their colors denote stability. The specific parameter values used are *c*_*h*_ = 0.25 and *d*_*hp*_ = 0. (B) Heatmaps of predicted equilibria and their stability over a range of additional host colonization caused by the mutualist and inherent mutualist colonization. Panels show different choices of the inherent host colonization rate. Here we assume that *d*_*hp*_ = 0. (C) Dynamics of the host, pathogen and mutualist over time when the feasible equilibrium is unstable (ie. the orange regime of Fig. 3B). (D) The fraction of the mutualist plotted against the fraction of the pathogen for different values of the additional host colonization due to the pathogen (panels). Color indicates time. The red point indicates the state of the system at the end of the simulation (at time *t* = 2000), which is not necessarily an equilibrium point. (E) Heatmaps displaying the coexistence outcomes (as a numerically stable limit cycle) of simulated dynamics (colors) as a function of the initial proportions of the host and the mutualist. Points where the fraction of the mutualist would exceed the fraction of the host are not included (white area). The parameter values are *c*_*h*_ = 0.5, *c*_*hm*_ = 10, *d*_*hp*_ = 0, and *c*_*m*_ = 10 with the remaining parameter values given in Table A1.

Further simulations reveal limit cycles when the equilibrium is feasible but unstable (Fig. 3C), emerging abruptly as we change the benefit of mutualists to hosts (Fig. 3D). These dynamics come about because high densities of each of the three species favor the growth of a different species in a cyclical fashion – a phenomena we describe in more detail in the Discussion. Systems with additional host colonization from the mutualist (*c*_*hm*_ > *c*_*h*_) also exhibit alternative stable states or bistability: depending on the starting conditions, populations will either collapse or exhibit long-term coexistence (Fig. 3A). Specifically, the initial proportions of the host and mutualist must be sufficiently high to ensure that the three species coexist (Fig. 3E). Obligate hosts require a threshold density of mutualists for population growth, and similarly, the mutualists need enough hosts to colonize. This reciprocal positive feedback between hosts and mutualists produces bistability. **Our third main result is that host colonization induced by the mutualist promotes coexistence even when the host populations cannot grow alone, but this coexistence may be dynamically complex**.

## Discussion

Our findings indicate that coexistence states – whether stable or unstable, fixed or periodic – are tightly coupled to symbiont effects on the host. Recent empirical work (Wang et al., 2022; Krediet et al., 2013) supports the basic structure of our model by finding similar dynamics among mutualistic and pathogenic bacteria and their coral hosts. In the experiments of Wang et al. 2022, pathogenic bacteria use hydrogen peroxide to induce lysis in temperate phages within mutualistic bacteria, killing these cells. Through this process, bacteria that are pathogenic to coral may obtain a competitive advantage over faster-growing and mutualistic bacteria. However, the effects of pathogenic bacteria may reduce coral growth rates and prevent stable coexistence in this system, eventually leading to the exclusion of the pathogen if mortality is too high. Our model results provide theoretical guidance for this experimental system, illustrating how feedbacks between microbial interactions and host dynamics can influence community stability. This insight extends beyond coral, with implications for other host-microbe systems. In particular, our findings are relevant to plant-soil feedback theory, as microbial communities colonizing plant surfaces and root systems exhibit similar dynamics (van der Putten et al., 2013). More broadly, the main takeaway from our work – that the structure of the holobiont can modify our classic understanding of coexistence – is quite general. This result has wide-ranging applications across various ecological systems where host-microbe interactions play a crucial role.

We modified the standard competition-colonization model by incorporating a dynamic “landscape” in the form of a coral host whose vital rates may be modified by bacterial colonization. Including a dynamic host – without any direct effect of bacteria on coral life history – sometimes counterintuitively altered the standard coexistence criteria. At the scale of an individual host, host death is unambiguously detrimental for both bacterial species. Yet, in the context of these competitive dynamics, increased host death benefits the mutualist over the pathogen. How does this difference come about? Intuitively, the superior competitor or pathogen is most limited by dispersal, and so the loss of available habitat is particularly detrimental to its ability to persist. The inferior competitor or mutualist disperses more rapidly and therefore benefits when the host undergoes fast turnover. The asymmetry between pathogen and mutualist broadens the regime in which these bacterial species coexist compared to the usual competition-colonization trade-off (Fig. 1B). This outcome has been previously observed in competition-colonization models in which habitat destruction is exogenously imposed (Hastings, 1980; Tilman, 1994; Yu and Wilson, 2001; Nee and May, 1992; Tilman et al., 1994, 1997; Amarasekare, 2003).

The cycles that occurred when bacteria were mutualistic highlighted the complex interplay between host, mutualist and pathogen dynamics. A closer examination of the trajectories in Fig. 3C helps explain this phenomenon. First, mutualists increase the fraction of unoccupied hosts because they boost the host colonization rate. The mutualists then take over these new hosts (unoccupied sites) because they are faster dispersers than the pathogens. The proportion of pathogens eventually increases, reducing the number of mutualists because the pathogenic bacteria are the dominant competitors. As a result, the proportion of hosts decreases, because the hosts are poor dispersers without the mutualists. Finally, the fraction of pathogens drops precipitously because it is especially sensitive to low host densities (as we saw in our first set of results). At this point, the cycle can begin again, with the growth of unoccupied hosts leading to an increase in the fraction of mutualists. Similarly, mutualistic bacteria produced another new dynamical property of our model – alternative stable states (Scheffer et al., 2001; Scheffer, 1989; Staver et al., 2011). Interestingly, alternative stable states in coral cover have been experimentally and theoretically demonstrated before (Mumby et al., 2007; Blackwood et al., 2018; Schmitt et al., 2019), though through a different mechanism, namely competition for space between coral and macroalgae (Mumby et al., 2007). These findings support previous research that the composition of bacterial communities may play a crucial role in coral ecosystem dynamics (Reshef et al., 2006). Further efforts are needed to explore how this understanding could potentially inform coral restoration efforts, particularly in terms of optimizing bacterial community composition in outplanted corals to promote the formation of stable holobionts.

In the model introduced here, prophage induction is simply the mechanism by which the pathogen kills the mutualistic bacteria. Empirical work by Wang et al. 2022 shows that the pathogen *V. coralliilyticus* releases hydrogen peroxide, which triggers the lytic cycle of a prophage present in the mutualist’s genome. Our model, however, does not capture the population dynamics of this temperate phage and its host. The decision between the lytic and lysogenic lifecycles by temperate phages has been the focus of theoretical work examining the optimal strategy under changing host and environmental conditions (Santillán and Mackey, 2004; Anthenelli et al., 2020; Cheong et al., 2022). Recent work has proposed a fitness switch, where the lysogeny is favored when the phage population grows faster as prophage than as virions produced by lysis, and conversely for the switch from lysogenic to lytic during induction (Roughgarden, 2024). Empirical work shows that lytic activity is significantly associated with the amount of live coral on Pacific coral reefs (Silveira et al., 2023). Therefore, it is possible that this additional layer of population dynamics between temperate phages and their bacterial hosts may affect the coexistence dynamics between bacteria and corals.

Throughout our work, we have only considered ordinary differential equations without stochasticity, but future work could include more mathematical complexity. Biologically, we modeled horizontal transmission – acquisition of microbes from the environment rather than passed down from parent to offspring. While this is appropriate for most coral species, some species exhibit vertical transmission in a strategy known as brooding (Leite et al., 2017; Damjanovic et al., 2020). Extending this model to include vertical transmission may be fruitful in further understanding the dynamics between hosts and microbes, particularly in the coral holobiont system. Many previous empirical studies of coral systems focus on coexistence outcomes when incorporating various species interactions such as competition with algae for space (Roach et al., 2020) or predation by parrotfish (Ezzat et al., 2020). Similarly, theoretical work has analyzed how temperature changes (Lima et al., 2020) or external perturbations (Dunphy et al., 2021) shift the structure of microbial communities on coral reefs. As a result, one of the most promising future directions in the theory of holobionts is to determine how the interaction between internal microbiome ecology and external community dynamics affects the ecosystem as a whole (Bell and Bordenstein, 2022).

## Acknowledgements

We thank Jie Deng, Pamela Ferreti, Armun Liaghat, Maria Martignoni, Taom Sakal, Athmanathan Senthilnathan, Lucas Souza, Chuliang Song, Bethany Stevens, Annie Innes-Gold, Sophia Rahnke, Elizabeth Gross, Seth Bordenstein, Zoe Cardon and Joan Roughgarden for insightful comments, discussion and feedback as part of the the Theory of Microbial Symbiosis Workshop at the Hawai’i Institute for Marine Biology. The authors gratefully acknowledge support from the Gordon and Betty Moore Foundation (GBMF10000) to attend the Theory of Microbial Symbiosis Workshop. TLG was supported by the National Science Foundation Graduate Research Fellowship Program under Grant No. DGE-2039656. KJMD was supported by the National Science Foundation under Grant No. MPS-2316455. JB was supported by the National Science Foundation Graduate Research Fellowship program under Grant No. DGE-2439024. LCM was supported by the National Science Foundation under Grant No. DBI-2233983.

## Statement of Authorship

TLG, JB and KJMD conceived of the project, performed mathematical analyses and wrote the first draft. TLG and KJMD wrote code for simulations. All authors interpreted model results and contributed substantially to editing the manuscript.

## Data and Code Availability

All the code used to generate figures is available at https://github.com/theogibbs/HostCompCol. There is no other data associated with this study.

## Appendix A: Additional Methods and Parameters

Table A1 displays the default values and ranges of the model parameters that we explored in simulations. The following sections present calculations of when the equilibria of our model are feasible and stable in the case when the bacteria do not affect the host.

**Table A1.** The model parameters with associated descriptions, baseline values and ranges, and units.

| Symbol | Description | Value | Range | Units |
| --- | --- | --- | --- | --- |
| $c_h$ | coral colonization rate | 2 | [0.5, 10] | per time |
| $c_{hm}$ | coral colonization rate when colonized by the mutualist | $c_h$ | [1.5, 10] | coral per mutualist per time |
| $c_p$ | pathogen colonization rate | 5 | | per coral per time |
| $c_m$ | mutualist colonization rate | 6 | [0, 16] | per coral per time |
| $d_h$ | coral mortality rate | 1 | [0, 1.5] | per time |
| $d_p$ | pathogen mortality rate | 1 | | per time |
| $d_{hp}$ | additional mortality from pathogenicity | 0 | [0, 2] | per time |
| $d_m$ | mutualist mortality rate | 1 | | per time |

## Feasibility criteria

Any equilibrium of (1) where all three species have positive abundance must satisfy the equations

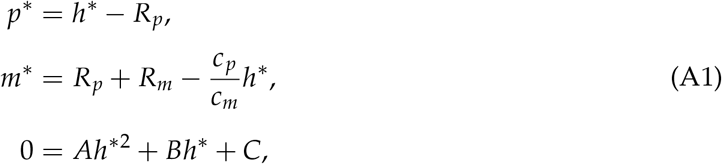

where 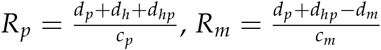 and

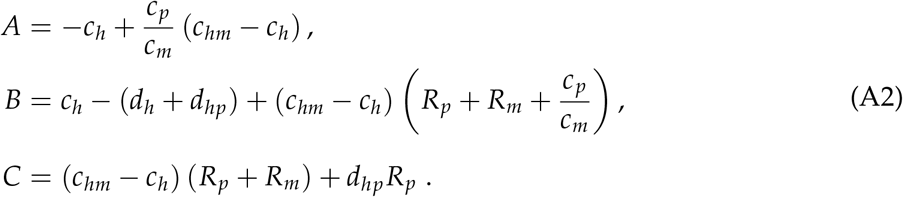

To determine feasibility in the main text, we solved these equations numerically and checked the signs of their solutions.

## No effects on host

Consider the situation where the bacteria do not impact the host: *c*_*hm*_ = *c*_*h*_ and *d*_*hp*_ = 0. In this scenario, *a* = −*c*_*h*_, *b* = *c*_*h*_ − *d*_*h*_ and *c* = 0 so that 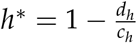 if it is non-zero. We find that

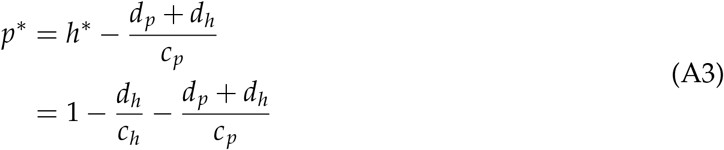

which is positive whe

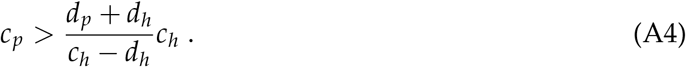

As *d*_*h*_ → 0, this inequality reduces to *c*_*p*_ > *d*_*p*_ as we expect. On the other hand, increasing *d*_*h*_ from below *c*_*h*_ always makes this inequality harder to satisfy, as we found in the main text. We also find

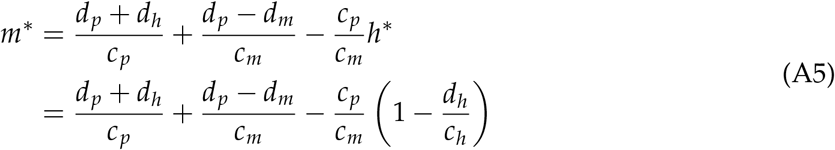

which is positive when

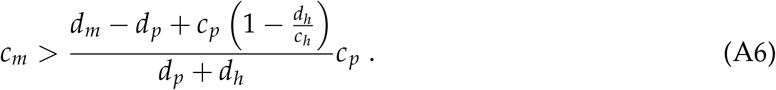

Once again, as *d*_*h*_ → 0, we get 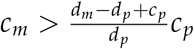 which is the standard formula for the competition-colonization tradeoff (Yu and Wilson, 2001). In this case, however, increasing *d*_*h*_ actually increases the proportion of mutualists at equilibrium, unlike in the case of the pathogen.

## Pathogen-induced host mortality

Now, we still take *c*_*hm*_ = *c*_*h*_, but we let *d*_*hp*_ > 0. Because the host dynamics now depend on the pathogen, its equilibrium fraction is given by a quadratic whose solution is

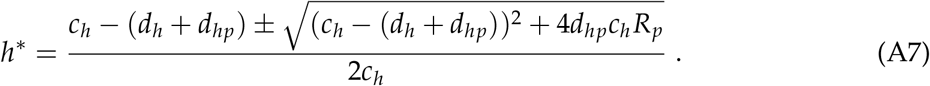

By assumption, 4*d*_*hp*_*c*_*h*_*R*_*p*_ > 0, so we know that the term in the square root is larger in absolute value than |*c*_*h*_ − (*d*_*h*_ + *d*_*hp*_)|. As a result, there is at most one positive root for *h*^*^, which is given by (A7) when the plus-minus is a plus.

## Stability analysis

After some rewriting, the Jacobian of our model evaluated at any non-trivial and feasible fixed point (*h*^*^, *p*^*^, *m*^*^) can be written as

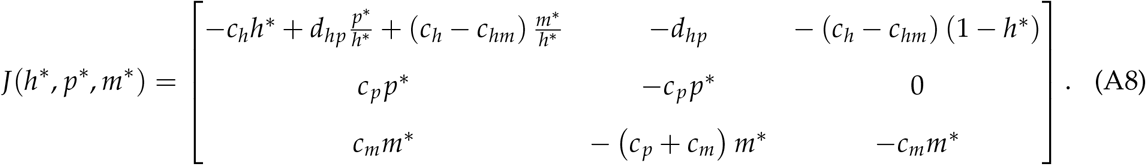

To determine stability in the main text, we evaluated this Jacobian at a given feasible equilibrium and then compute its eigenvalues numerically.

## No effects on host

When there is no effect of the bacteria on the host, the Jacobian simplifies to

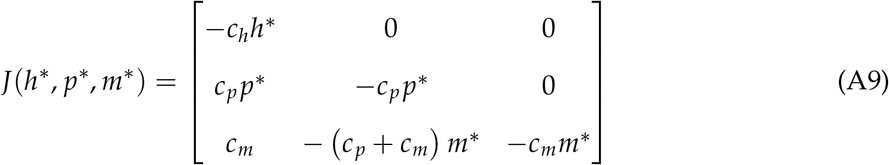

The eigenvalues of this lower-triangular matrix are simply

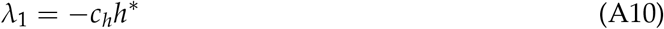

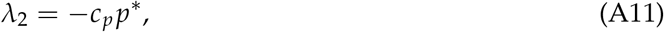

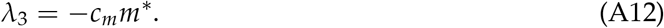

Therefore, in this case, if a fixed point is feasible it must also be stable.

## Notes

### Competing Interest Statement

The authors have declared no competing interest.

